# SEPIA - SuscEptibility mapping PIpeline tool for phAse images

**DOI:** 10.1101/2020.07.23.217042

**Authors:** Kwok-Shing Chan, José P. Marques

## Abstract

Quantitative susceptibility mapping (QSM) is a physics-driven computational technique that has a high sensitivity in quantifying iron deposition based on MRI phase images. Furthermore, it has a unique ability to distinguish paramagnetic and diamagnetic contributions such as haemorrhage and calcification based on image contrast. These properties have contributed to a growing interest to use QSM not only in research but also in clinical applications. However, it is challenging to obtain high quality susceptibility map because of its ill-posed nature, especially for researchers who have less experience with QSM and the optimisation of its pipeline. In this paper, we present an open-source processing pipeline tool called SuscEptibility mapping PIpeline tool for phAse images (SEPIA) dedicated to the post-processing of MRI phase images and QSM. SEPIA connects various QSM toolboxes freely available in the field to offer greater flexibility in QSM processing. It also provides an interactive graphical user interface to construct and execute a QSM processing pipeline, simplifying the workflow in QSM research. The extendable design of SEPIA also allows developers to deploy their methods in the framework, providing a platform for developers and researchers to share and utilise the state-of-the-art methods in QSM.

## 1. Introduction

Quantitative susceptibility mapping (QSM) provides a unique image contrast based on tissue magnetic susceptibility. Brain tissues containing paramagnetic substances that enhance local magnetic fields, such as irons stored in ferritin in the deep grey matter or hemosiderin in haemorrhages, can be easily distinguished from the tissues containing diamagnetic substances that weaken the local magnetic fields, including calcification or myelin in white matter. Unlike in conventional T_1_-weighted and T_2_*-weighted imaging, these two types of substances generate opposite image contrasts in QSM, making it particularly useful in applications where the tissues with opposite properties need to be distinguished, such as between haemorrhages and calcifications (Chen et al., 2013). Additionally, strong correlations were demonstrated in various *ex vivo* studies between iron concentration in the deep nuclei and magnetic susceptibility (Langkammer et al., 2012; Sun et al., 2015), showing evidence of the quantitative ability of QSM which can find applications on studying, for example, the relationships between iron overload and neurodegenerative diseases, including Parkinson’s disease (He et al., 2015; Ide et al., 2015; Langkammer et al., 2016; Mazzucchi et al., 2019; Thomas et al., 2020), Huntington’s disease (Chen et al., 2018; D et al., 2015), Alzheimer’s disease (Acosta-Cabronero et al., 2013; Bulk et al., 2020; Du et al., 2018; Gong et al., 2019; O’Callaghan et al., 2017), and between iron deposition in the brain and ageing (Acosta-Cabronero et al., 2016; Betts et al., 2016; Bilgic et al., 2012; Gong et al., 2015; Keuken et al., 2017; Liu et al., 2016; Persson et al., 2015).

In a nutshell, QSM relies on the measurement of magnetic field heterogeneity induced by brain tissues inside a strong magnetic field and the application of a dipole inversion to compute the magnetic susceptibility distribution, as illustrated in Fig. 1. The phase of a 3D GRE data is typically used to measure the field heterogeneity. However, the phase value of MRI data is limited by a range between -π and π, which originates phase wrapping artefacts. Before any further computation, spatial and temporal phase unwrapping are usually needed to obtain the true phase evolution (this is particularly important in regions close to large magnetic susceptibility sources with data acquired at long echo time). Subsequently, the unwrapped phase data acquired at a single or multiple time points is used to compute the total frequency deviations from the reference Lamour frequency, assuming linear phase accumulation over time (Robinson et al., 2017).

**Fig. 1:**
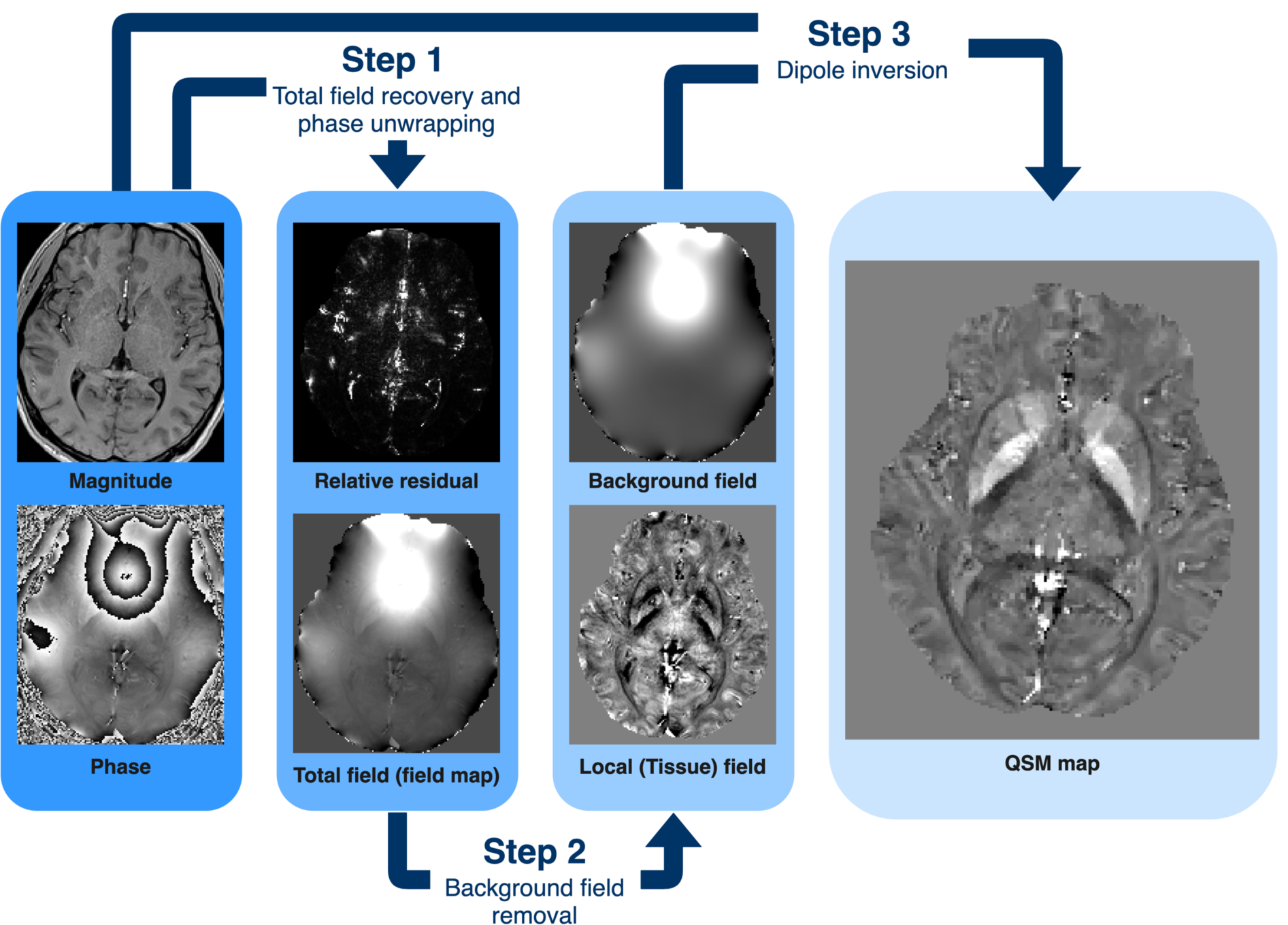
Summary of a standard QSM processing pipeline. MRI phase data has to be pre-processed to remove phase wrapping artefact and non-tissue field contributions before computation of QSM (from left to right). In most cases, each step of QSM processing is performed separately and therefore care has to be taken to ensure high quality intermediate results as error can propagate to the final QSM map computation. Magnitude part of the data is often used as prior information in the QSM dipole inversion to mitigate error because the process is ill-conditioned. The relative residual map derived from mono-exponential fitting can be used to further improve QSM processing results.

The field deviations measured in MRI are caused by the magnetic susceptibility both of the tissue of interest and the field perturbations originated from elsewhere (Li, 2001; Schweser et al., 2016a). The frequency components from non-tissue background sources, such as air/tissue interface, air/bone interface and scanner hardware imperfection, need to be removed from the total frequency map so that only the frequency components coming from the brain tissues remain in subsequent processing (Schweser et al., 2016b). As there are more potential sources of background field (both within and outside the field of view of the acquisitions) than detectors (data with signal within the FOV, but outside the region of interest) makes the mapping of the background sources extremely difficult and ill-posed. Fortunately, the magnetic fields originated from the background sources within the region of interest differ significantly from the brain tissue fields (Schweser et al., 2016b), which make the background field removal possible.

Finally, mapping susceptibility sources from the tissue field map can be done by deconvolution of the tissue field and a unit dipole field. This operation, or so-called dipole inversion, can be natively performed in the k-space domain where the Fourier transform of the image-space deconvolution is equivalent to the division of the Fourier transforms of the tissue field by the unit dipole. However, there is an infinite number of magnetic susceptibility distributions that can explain the measured field. Numerically, this is the result of the Fourier transform of the dipole field having a cone surface of zero values, making the division undefined. Furthermore, the vicinity of this conical surface generates significant noise amplification that can result in streaking artefacts in the final susceptibility map. To overcome this ill-posed inversion problem, various methods have been suggested over the last decade to incorporate various types of prior information such as k-space thresholding (Shmueli et al., 2009; Wharton et al., 2010), spatial smoothness of the susceptibility map (Bilgic et al., 2014a), consistence of detectable brain structures between the susceptibility map and magnitude data (J. Liu et al., 2012; Liu et al., 2011b, 2018), among many others more advanced methodologies introduced recently. For a more thorough review of QSM methods refers to (Deistung et al., 2016; Wang and Liu, 2015).

Recent developments in the QSM field have generated an invaluable set of resources online. Extensive methods and codes have been shared for academic purposes in the websites of various research groups and software development platforms such as GitHub, GitLab and MathWorks File Exchange. While MATLAB (Natick, USA) is the current main software environment for QSM development, codes developed in Python are also available. Efforts were also made to include QSM in MRI data processing frameworks such as brain imaging data structure (BIDS) apps, statistical parametric mapping (SPM) (https://www.fil.ion.ucl.ac.uk/spm/) and qMRLab (http://qmrlab.org/). Some of the QSM software packages can be found on a GitHub page (https://github.com/mathieuboudreau/qsm-tools).

Despite the vast resources available online making high quality QSM possible, the convoluted nature of the QSM processing can hinder its wider application in research and clinical settings, especially for researchers who want to apply the method in their studies but have less experience with the QSM processing and theory. Certain software packages such as MEDI toolbox and STI Suite provide a graphical user interface (GUI) to ease the use for data processing. However, these packages often support only established works within their research groups, limiting the ability to include third-party algorithms in one or several parts of the processing pipeline to achieve a better QSM result. Integrating parts of the different software packages is possible in command-line based operations, yet compatibility between toolboxes including data format and unit conversion is not always straightforward. Nevertheless, combining many of these methods is generally accepted to be beneficial and the optimum processing pipeline is very much dependent on the data available (resolution, SNR, multi-echo vs single echo) and will, therefore, be study-specific.

In this work, we present the SuscEptibility mapping PIpeline tool for phAse images (SEPIA) toolbox for QSM and other phase processing methods. In contrast to the existing software for QSM, SEPIA focuses on the integration of various QSM toolboxes in a unified platform, providing diverse options for researchers, thus maximising the possibility of computing high quality susceptibility maps, together with the flexibility of including new methods available in the future. SEPIA provides both a GUI to compose QSM processing pipeline intuitively and the option to script the resulting pipeline, either for optimisation purpose or to apply to a large cohort of patients.

## 2. Methods

### 2.1. Dependency, Installation and Documentation

SEPIA is a QSM processing pipeline tool developed in MATLAB providing with both graphical and command-based operations that allows easy access to various processing algorithms in a unified platform, both for less-experienced users and advanced researchers. The software is supported from MATLAB version R2014b onwards on operating systems including Linux, Mac and Windows. It is open-source software with the MIT license and can be downloaded from the GitHub page, https://github.com/kschan0214/sepia. Full documentation including installation, functionality and tutorial is available in https://sepia-documentation.readthedocs.io/.

The latest version of SEPIA (version 0.8.0) is optimised to access the following toolboxes and processing methods for QSM that are freely available for academic purposes, including the MEDI toolbox (updated 15 January 2020), STI suite (version 3.0), FANSI toolbox (commit dc68×306) and Speedy rEgion Growing for Unwrapping Estimated phase (SEGUE) (Karsa and Shmueli, 2018), along with other QSM and SWI method implementations (Abdul-Rahman et al., 2007; Bilgic et al., 2014a; Li et al., 2011; Polak et al., 2020; Schweser et al., 2011; Sun and Wilman, 2014; Wharton et al., 2010).

### 2.2. SEPIA architecture

SEPIA is designed with a GUI frontend and a data processing backend to achieve various QSM processing tasks with the design architecture illustrated in Fig. 2. The GUI provides an intuitive way to construct a QSM processing pipeline through the selection and combination of methods from various toolboxes and its functions will be discussed in the next section. The core of SEPIA is divided into three levels, namely the I/O level, the task level and the communication level. The I/O level handles user input data (images in the NIfTI format), including validation of the input and conversion to MATLAB variables that can be used in the task level. It also generates output maps in the NIfTI format with an identical geometric coordinate as the input data.

**Fig. 2:**
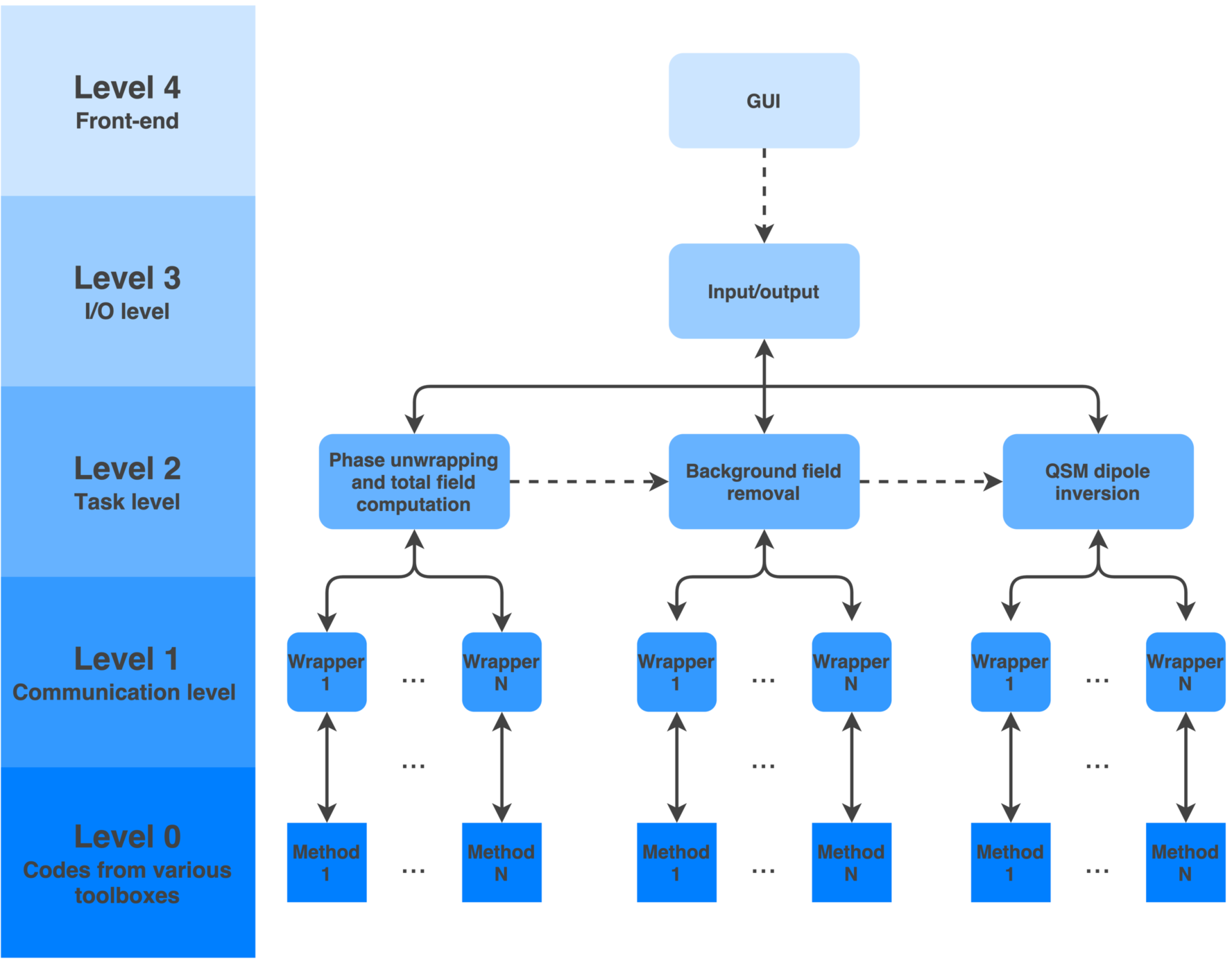
Schematic of SEPIA computational architecture. The design of SEPIA has a depth of four levels responsible for specific objectives. The top level is a GUI frontend providing easy access to all supported methods to construct a QSM processing pipeline. It creates a pipeline configuration file to be used in the backend where all the processing procedures happen (levels 0-3). The I/O level is a wrapper of the tasks to be performed in the task level, supporting input and output with NIfTI data format instead of MATLAB variables. Depending on the user choices, methods from various toolboxes will be called by SEPIA via the method wrapper function in the communication level to perform a designated task. Arrow direction indicates how information is communicated within the SEPIA framework.

In the task level, QSM data processing is divided into three primary tasks:

1. computation of total frequency shift from the (multi-echo) MRI phase data (see section 2.3.2. Total field recovery and phase unwrapping panel);
2. separation of the tissue field from the background field in the total frequency shift map (see section 2.3.3. Background field removal panel);
3. computation of magnetic susceptibility from the tissue field (see section 2.3.4. QSM panel). These tasks can be accomplished successively or separately, depending on the user specification and interface used (see Fig. 3). The algorithm selected by the user, which is categorised based on the tasks, will then be called in the task level.

The communication level is the connection between SEPIA and the methods provided in the supported toolboxes. It contains wrapper functions that parse the user-defined algorithm parameters and obtain required input data in the SEPIA structure to the format that is required by the selected method. With the wrapper functions in the communication level, QSM method developers do not need to modify their existing methods in order to add them to the SEPIA framework, giving the flexibility to incorporate the latest QSM methods in the future.

**Fig. 3:**
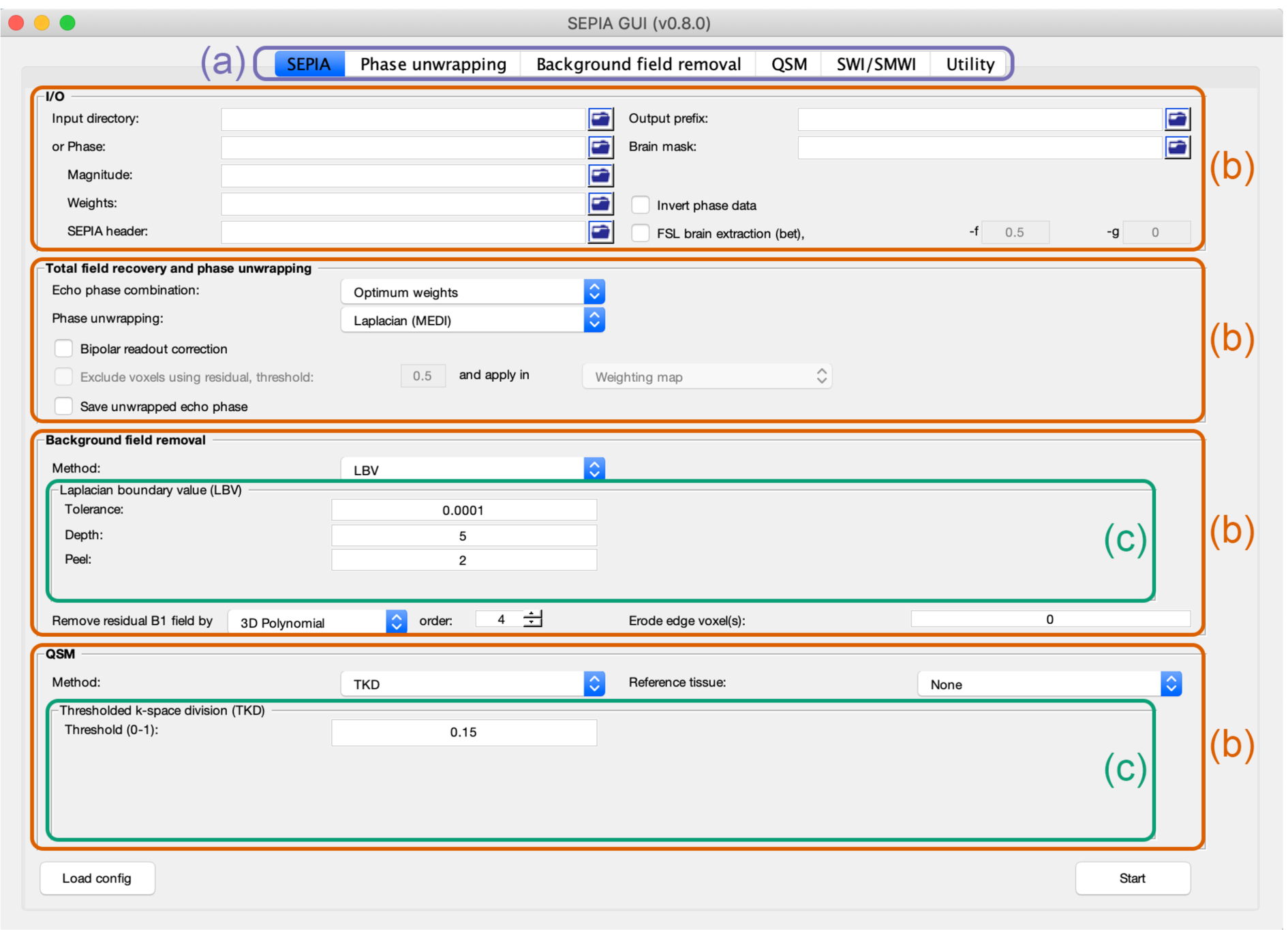
An illustration of the GUI of SEPIA. The functionality of the GUI is divided into three sections. (a) Users can switch between various applications using the tabs on the top of the GUI to perform a specific processing pipeline as their names suggested. (b) Each application consists of multiple panels designed for a specific task, including data input/output, total field recovery, background field removal and QSM dipole inversion. (c) For the methods in background field removal and QSM dipole inversion steps, each of them has its own method panel allowing users to fine-tune the results of these steps.

### 2.3. Graphical user interface (GUI) and functionality

The GUI of SEPIA provides the most straightforward way to access the full functionality of the software (Fig.3). All supported algorithms are available as selectable options in the GUI. Users can mix and match methods from different toolboxes to build a processing pipeline optimised for their needs without paying extra time on adapting data from one software package to another. Instead, users are encouraged to focus on exploring combinations of different methods and fine-tuning algorithm parameters such as regularisation value to achieve the best results for their needs, as the ill-posedness of QSM processing together with the variety of imaging protocols and related SNR or artefacts limit the generality of a universal pipeline application and often returns sub-optimal results.

The GUI consists of various ‘tabs’ corresponding to various applications tackling different tasks in the phase data processing (Fig. 3a). Currently available tabs include:

- **SEPIA**: construction of a full QSM processing pipeline from raw phase data to magnetic susceptibility map;
- **Phase unwrapping**: construction of a pipeline to compute a total field map (frequency shift) from raw phase data;
- **Background field removal**: construction of a pipeline to compute a (local) tissue field map from a total field map;
- **QSM**: construction of a pipeline to compute a magnetic susceptibility map from a tissue field map;
- **SWI/SMWI**: construction of a pipeline to perform susceptibility-weighted imaging (SWI) (Haacke et al., 2004; Reichenbach et al., 1997) or susceptibility map-weighted imaging (SMWI) (Gho et al., 2014) from complex-valued MRI data; and
- **Utility**: containing functions that can assist QSM processing.

The QSM operations in the GUI are divided into 4 major panels aiming at specific tasks individually, namely (1) I/O panel, (2) Total field recovery and phase unwrapping panel, (3) Background field removal panel and (4) QSM panel.

#### 2.3.1. I/O panel

The I/O panel allows users to specify the input and the output. Typically, the phase and magnitude MRI NIfTI images and a signal mask are required to compute a QSM map. The input images should be in 3D (single echo) or 4D (multi-echo with time in the last dimension) NIfTI format. The data can be acquired with a 2D or 3D sequence with any arbitrary readout strategies as long as the phase images are available. The magnitude input has inherently arbitrary units, while the phase input can be in radian unit (typically [-π, π]) or DICOM values (int12 - [0, 4096] or [-4096, 4096]), which will be rescaled to radian.

To be able to compute a frequency shift map and a susceptibility map with correct units, it is important to inform the corresponding algorithms with additional information including the main magnetic field (B_0_) strength, B_0_ direction with respect to the imaging field of view and echo times of the acquisition. This information is not usually available in the NIfTI header but is accessible in supplementary files in text or JavaScript Object Notation (JSON) formats, depending on the DICOM conversion tools. SEPIA has a function to extract the information from the supplementary files and to store it in a specific MATLAB format file (.mat) which is part of the essential input for SEPIA called “SEPIA header” (see “Utility” panel).

Two input routines are supported in SEPIA: users can select an input directory which contains all the essential files with a pre-defined naming structure (see Supplementary Table S1) or the required files can be manually inserted in the corresponding fields. As an alternative to provide a signal mask, it is possible to perform brain extraction on whole-brain magnitude data based on the MATLAB implementation of the *BET* tool of FSL from the MEDI toolbox. The full list of the SEPIA outputs and their description are provided in Table 1.

**Table 1:** Full list of filenames of SEPIA outputs.

| Phase unwrapping application output directory | Description |
| --- | --- |
| <outputPrefix>_total-field.nii.gz | Unwrapped total frequency shift map, in Hz. |
| <outputPrefix>_weights.nii.gz | SNR-weighted image derived from standard deviation of noise in phase data. See Supplementary Eq. S2. |
| <outputPrefix>_noise-sd.nii.gz | Estimated standard deviation of noise in the phase data. See Supplementary Eq. S1. |
| <outputPrefix>_phase.nii.gz | Wrapped phase data in radian (only if the input data contains voxel exceeds the range of $[-\pi, \pi]$). |
| <outputPrefix>_unwrapped-phase.nii.gz | Unwrapped phase data in radian (if selected). |
| <outputPrefix>_mask.nii.gz | Brain mask derived from *BET* of FSL (if selected). |
| <outputPrefix>_mask-local_field.nii.gz | Signal mask for background field removal step (if voxel exclusion is selected with 'Brain mask' option ). |
| <outputPrefix>_relative-residual.nii.gz | Relative residual derived using a mono-exponential model with a single frequency shift (if voxel exclusion is selected). See Supplementary Eqs. S3-4. |
| <outputPrefix>_mask-reliable.nii.gz | Derived from thresholding the relative-residual map using user-defined value. |
| Background field removal application output directory | Description |
| <outputPrefix>_local-field.nii.gz | Local (tissue) field map, in Hz. |
| <outputPrefix>_mask-qsm.nii.gz | Signal mask for QSM step. |
| QSM application output directory | Description |
| <outputPrefix>_QSM.nii.gz | Magnetic susceptibility map, in ppm. |
| Other output files | Description |
| sepia_config.m | Script automatically generated by the GUI of SEPIA containing all user specified parameters. |
| run_sepia.log | Event log file of the MATLAB's command window output. |
| run_sepia.error | Error message of SEPIA (if any). |
| SWI/SMWI application output directory | Description |
| <outputPrefix>_swi-phase.nii.gz | High-pass filtered phase data. |
| `<outputPrefix>_swi-positive.nii.gz` | Positive phase susceptibility-weighted images (if selected). |
| `<outputPrefix>_swi-mIP-positive.nii.gz` | Positive phase susceptibility-weighted images with minimum intensity projection over user-defined field of view (if selected). |
| `<outputPrefix>_swi-negative.nii.gz` | Negative phase susceptibility-weighted images (if selected). |
| `<outputPrefix>_swi-mIP-negative.nii.gz` | Negative phase susceptibility-weighted images with minimum intensity projection over user-defined field of view (if selected). |
| `<outputPrefix>_smwi-paramagnetic.nii.gz` | Paramagnetic susceptibility map-weighted images (if selected). |
| `<outputPrefix>_smwi-mIP-paramagnetic.nii.gz` | Paramagnetic susceptibility map-weighted images with minimum intensity projection over user-defined field of view (if selected). |
| `<outputPrefix>_smwi-diamagnetic.nii.gz` | Diamagnetic susceptibility map-weighted images (if selected). |
| `<outputPrefix>_smwi-mIP-diamagnetic.nii.gz` | Diamagnetic susceptibility map-weighted images with minimum intensity projection over user-defined field of view (if selected). |

#### 2.3.2. Total field recovery and phase unwrapping panel

The total field recovery and phase unwrapping panel provides options to recover the actual frequency shift from wrapped phase data. There are six implementations available for phase unwrapping of the DICOM converted phase images, including two Laplacian-based methods from MEDI toolbox and STI Suite (Li et al., 2011; Schofield and Zhu, 2003), a region growing methods (Witoszynskyj et al., 2009), a 3D best path method (Abdul-Rahman et al., 2007), SEGUE (Karsa and Shmueli, 2018) and a graph-cut method (Dong et al., 2015). Two methods are supported to estimate the total field when the input has multiple echoes, including a nonlinear data fitting approach (T. Liu et al., 2012) and a weighted average method based on the SNR of the magnitude MRI data (Robinson et al., 2017). The unit of the output total field map is in Hertz (Hz).

When a bipolar readout gradient is used in multi-echo data acquisition, eddy current can be induced on the gradient coils, which can introduce an undesirable phase ramp between the odd-number and even-number echoes along the readout encoding direction. Such frequency shifts, that are not static over time, degrade the accuracy of the total field measurement as well as any metric that assumes a linear phase increment as a function of echo time, resulting in sub-optimal QSM reconstructions (Li et al., 2015). In the SEPIA framework, an option is provided to correct the phase offset by removing a 1^st^ order polynomial field from the phase data if the input has more than 4 echoes.

In QSM processing, errors in the measurement of the local field associated artefacts (such as flow) or low SNR can create strong steaking artefacts in the computed magnetic susceptibility map. SEPIA provides an option (see Fig. 3) to exclude those image voxels from subsequent computation, using the relative measurement error derived from fitting the data to a mono-exponential decay model with a single frequency shift (see Eqs. S3-4 in Supplementary material). This information can be either added to the signal mask or a weighting map which is used to weight the data consistency term and the prior information term in some QSM algorithms.

#### 2.3.3. Background field removal panel

Before computation of the magnetic susceptibility from the field map, magnetic fields contributed from external sources other than brain tissue, such as air/tissue interface, have to be removed from the tissue field on the total field map. There are six methods available in SEPIA: (1) Laplacian boundary value approach (Zhou et al., 2014), (2) projection onto dipole field (Liu et al., 2011a), (3) sophisticated harmonic artefact reduction for phase data (SHARP) (Schweser et al., 2011), (4) regularisation-enabled SHARP (RESHARP) (Sun and Wilman, 2014), (5) variable kernel SHARP (V-SHARP) (Li et al., 2011) and iHARPERELLA (Li et al., 2014). Adjustable algorithm parameters of each method can be directly accessed on the corresponding method panel on the GUI (Fig. 3b). The input and output data of this task are in Hz.

In the case of single echo data or methods that assume the initial phase (at TE=0) to be zero, phase contributions from transmit field by an RF coil (B1+) and B1- of a body coil can remain in the resulting tissue field map even after the removal of the background field. These phase contributions and any other residual background field can be reduced using SEPIA by removing an N-order (N ranged from 1 to 4) polynomial fields or spherical harmonic fields from the tissue field.

#### 2.3.4. QSM panel

The last step in QSM processing is to map the magnetic susceptibility sources based on the tissue field computed after the fields generated by external sources are removed, which is also known as dipole inversion. This task can be done on k-space or image space and a variety of methods is available in the QSM panel: (1) a thresholded k-space method (Shmueli et al., 2009; Wharton et al., 2010), (2) a closed-form solution with an L2-regularisation method (Bilgic et al., 2014a), (3) an iterative LSQR approach (Li et al., 2011), (4) Star-QSM (Wei et al., 2017, 2016, 2015), (5) MEDI+0 (J. Liu et al., 2012; Liu et al., 2011b, 2018), (6) FANSI with the weak-harmonic regularisation (Bilgic et al., 2015, 2014b; Milovic et al., 2019, 2018) and (7) a non-linear dipole inversion (NDI) method (Polak et al., 2020). Some algorithms utilise prior information such as magnitude images to constraint the resulting susceptibility map. In most instances, SEPIA uses an estimation of the noise standard deviation in the phase data or the magnitude images as the constraint. It is also possible for users to provide an alternative image (registered to the field data and with the same matrix size) of their preference as the prior for the dipole inversion process. The output of the magnetic susceptibility map is in parts per million (ppm).

To be able to compare susceptibility values either across groups or in longitudinal studies, it is important to define a reference region. This is of great importance because the derived susceptibility map is a relative measurement. In SEPIA, users can choose the mean susceptibility value of the whole brain or the cerebrospinal fluid (CSF) in the lateral ventricle (Liu et al., 2018) as the reference (in which case this region will have mean susceptibility 0).

#### 2.3.5. SWI and SMWI standalone application

In addition to QSM processing, SEPIA also supports contrast enhancement techniques using both magnitude and phase of MRI data, including SWI (Haacke et al., 2009) and SMWI (Gho et al., 2014). The SWI implementation in SEPIA is based on high-pass homodyne filtering (Noll et al., 1991), through the complex division between the input phase data and its low-pass filtered copy processed with a 2D Hamming filter. The kernel size of the Hamming filter is a tunable parameter in the GUI, which has a direct influence on the residual of the low-spatial frequency background field in the high-pass filtered phase and is useful to handle data at long echo times where phase wrapping is more frequent. The SWI result can be further optimised via the “Threshold (rad)” and “Contrast” options in the GUI, corresponding to the threshold value to suppress positive/negative phase in the weighting map, and the number of times this map to be applied to the magnitude data (Haacke et al., 2009). The outputs of the application include the high-pass filtered phase data and the SWI images. Optionally, users can also export the SWI images with minimum intensity projection along slice direction over a small slab which often uses to improve visualisation of venous structures.

SMWI is a contrast enhancement method similar to SWI (Gho et al., 2014). Nevertheless, it benefits from a better contrast localisation of QSM to avoid the blooming artefact present in phase images and incomplete phase wrapping removal after high-pass filtering. The SMWI application in SEPIA accepts magnitude data together with a susceptibility map (ideally from the output of SEPIA which has units in ppm) as input. The operations of SMWI is very similar to the SWI counterpart, where the degree of contrast enhancement is controlled by the “Threshold (ppm)” and “Contrast” options in the GUI. The outputs are the paramagnetic/diamagnetic enhanced SMWI images.

### 2.4. Pipeline configuration file and batch processing

The main goal of the GUI is to generate a pipeline configuration file containing all the tasks, methods and algorithm parameters specified by the users, which is used to trigger the QSM processing (Fig. 4). The configuration files automatically generated by the SEPIA GUI correspond to an executable MATLAB script to obtain the same result without initialising the GUI since it directly accesses the processing backend. Depending on the task involved in the processing, the I/O level function *sepiaIO* will automatically sort out the corresponding pipeline function based on the main input variable *algorParam* which is a structure that describes all the pipeline steps to be performed. The selected methods and the tunable parameters of each step are clearly specified in its sub-structures: “*unwrap*”, “*bfr*” and “*qsm*”. The full structure of *algorParam* is provided in Table S2 in the Supplementary material for users who are interested in using the command-based feature of SEPIA.

**Fig. 4:**
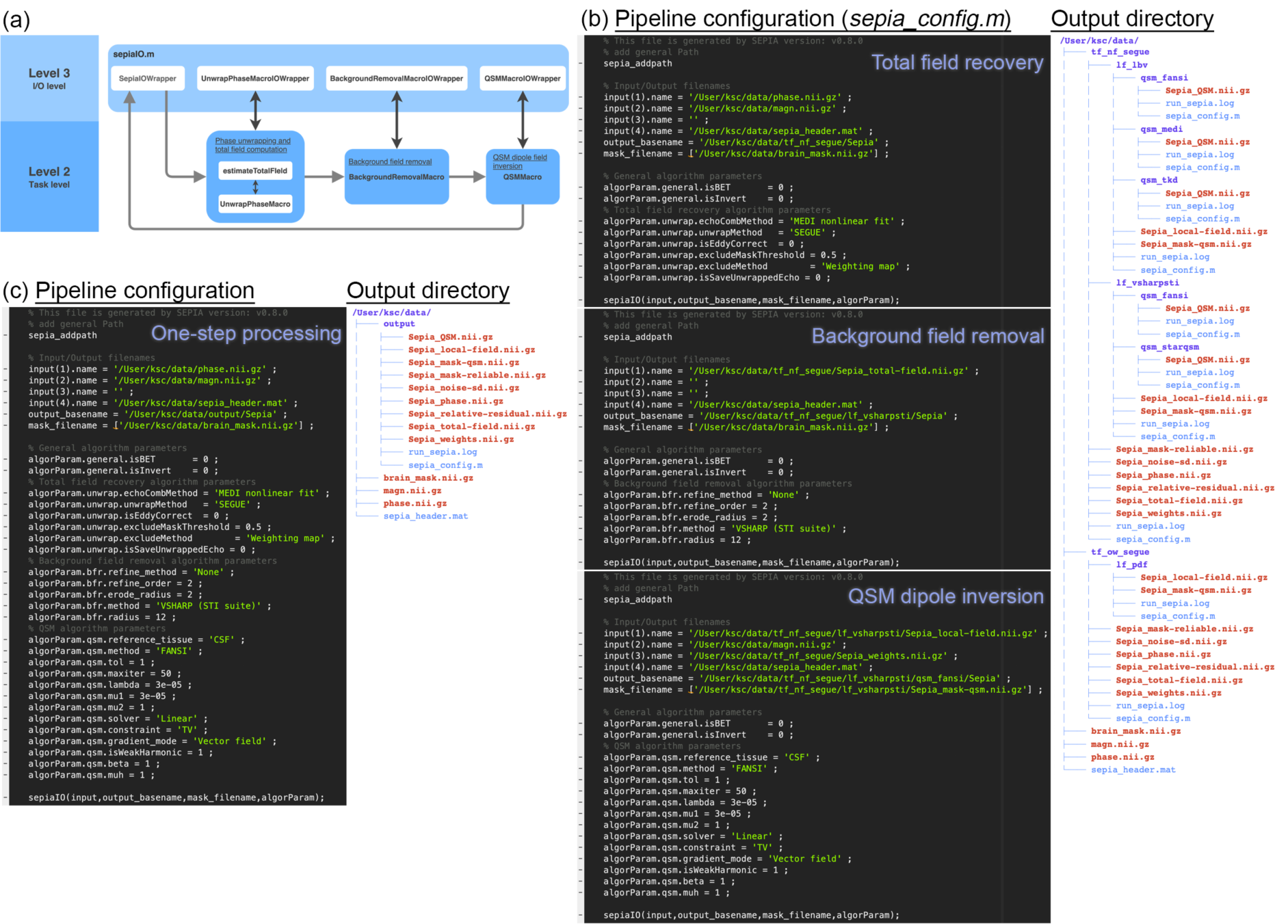
(a) Depending on the application selected, a corresponding wrapper function in the I/O level will be called to execute the task(s). (b) Typically, multiple methods or algorithm settings will be tested individually when optimising a QSM processing pipeline (right). In this example, a cascade of directories is created when the steps were run separately, allowing a complete track of the processing pipeline as each output directory contains a pipeline configuration file (*sepia_config*.*m*) specific to the method/parameters used to produce the result (left). (c) Once users have decided their processing pipeline, the pipeline configuration file of each step can be imported back to the SEPIA GUI and combined to create a single pipeline configuration file for a complete QSM processing. If the one-step application is used, the output directory will contain all output data from the separate steps (total field computation, background field removal and QSM calculation).

Importing the configuration file to the GUI is viable using the designated push button. This can be useful to re-use an existing pipeline from one study for a new dataset, or to finalise a full pipeline by concatenating multiple configuration files after the optimisation of individual processing tasks, such as processing method, regularisation value, iteration number or degree of brain erosion when performing background field removal (Fig. 4). The configuration file can also be useful for studies that involve a large number of datasets. Users can optimise the processing pipeline on a few datasets and then apply the same pipeline to the rest by making simple modifications in the configuration file such as adapting the filenames or directory accommodating the data. This process can even be more efficient if users have access to multiple processing units, such as a high-performance computing service, where batch processing can be easily deployed by running the configuration files on separate computing cores.

### 2.5. Integration of new algorithms to SEPIA

New phase unwrapping, background field removal and dipole inversion methods are constantly being developed to achieve further improvements in computational speed, image quality, robustness and/or accuracy of QSM values. SEPIA can serve as a framework to host the latest methods. New methods developed in the MATLAB environment can be integrated into the SEPIA processing backend smoothly via a wrapper function in the communication level as a connector. All the imaging data and algorithm parameters provided by the user can be accessed using this wrapper function, including the original complex-valued MRI image, acquisition information and resulting maps derived from previous processing steps with unified formats and units. Therefore, developers can adopt this comprehensive information without modification of their existing methods. Researchers can therefore develop new methods in their chosen units and function format as they preferred and can integrate the method retrospectively to SEPIA.

Additionally, if developers are interested to have their methods also shown in the GUI, they can design a graphical method panel to obtain the required algorithm parameters from user input. Detailed tutorial of how to integrate a new method to the SEPIA framework is also provided in the documentation website.

### 2.6. Examples of SEPIA usage

To demonstrate the flexibility of SEPIA on QSM processing pipeline creation, a test dataset, downloaded from the Quantitative Susceptibility Mapping of the Cornell MRI Research webpage (http://pre.weill.cornell.edu/mri/pages/qsm.html), was used to process with methods in various software packages supported in SEPIA. The test dataset was acquired on a 3T scanner (Verio, Siemens, Erlangen) with the following acquisition parameters:

Monopolar 3D multi-echo GRE with 8 echoes, TR/TE1/ΔTE/TE8=55/3.6/5.9/45ms, flip angle of 15°, resolution of 0.94 × 0.94 × 2 mm^3^, receiver bandwidth of 241 Hz/pixel, in-plane acceleration of 2 and partial Fourier of 6/8.

Three pipelines were built to compute QSM maps from the test dataset using the following methods:

1. Total field recovery: non-linear fitting (T. Liu et al., 2012) with Laplacian phase unwrapping (Schofield and Zhu, 2003); Background field removal: projection onto dipole field (Liu et al., 2011a); QSM dipole inversion: FANSI with weak-harmonic regularisation (Milovic et al., 2019).
2. Total field recovery: non-linear fitting (T. Liu et al., 2012) with graph-cut phase unwrapping (Dong et al., 2015); Background field removal: V-SHARP (Li et al., 2011); QSM dipole inversion: MEDI+0 (Liu et al., 2018).
3. Total field recovery: echo phase combination with optimum weights (Robinson et al., 2017) and SEGUE phase unwrapping (Karsa and Shmueli, 2018); Background field removal: Laplacian boundary values (Zhou et al., 2014); QSM dipole inversion: STAR-QSM (Wei et al., 2015).

Full details of the processing pipelines, including the specific algorithm parameters, is also illustrated in Table 2. Finally, to demonstrate the ability of SEPIA to the creation of both SWI and SMWI images, the original complex-valued data was processed with the default SWI method (threshold=π and contrast=4), and the QSM map derived from pipeline 3 was used in SMWI processing in SEPIA (threshold=1 ppm and contrast=4).

**Table 2:**
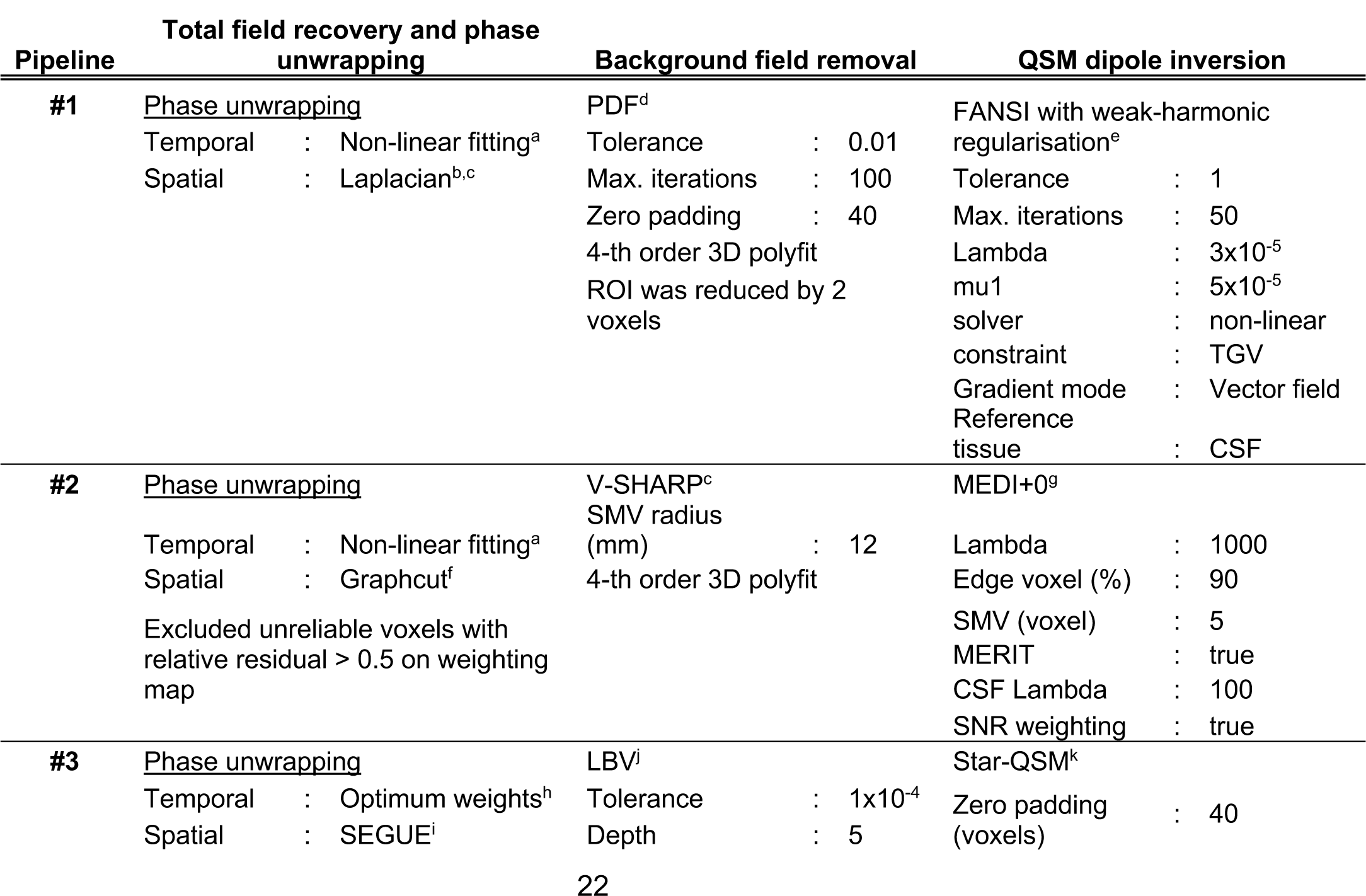
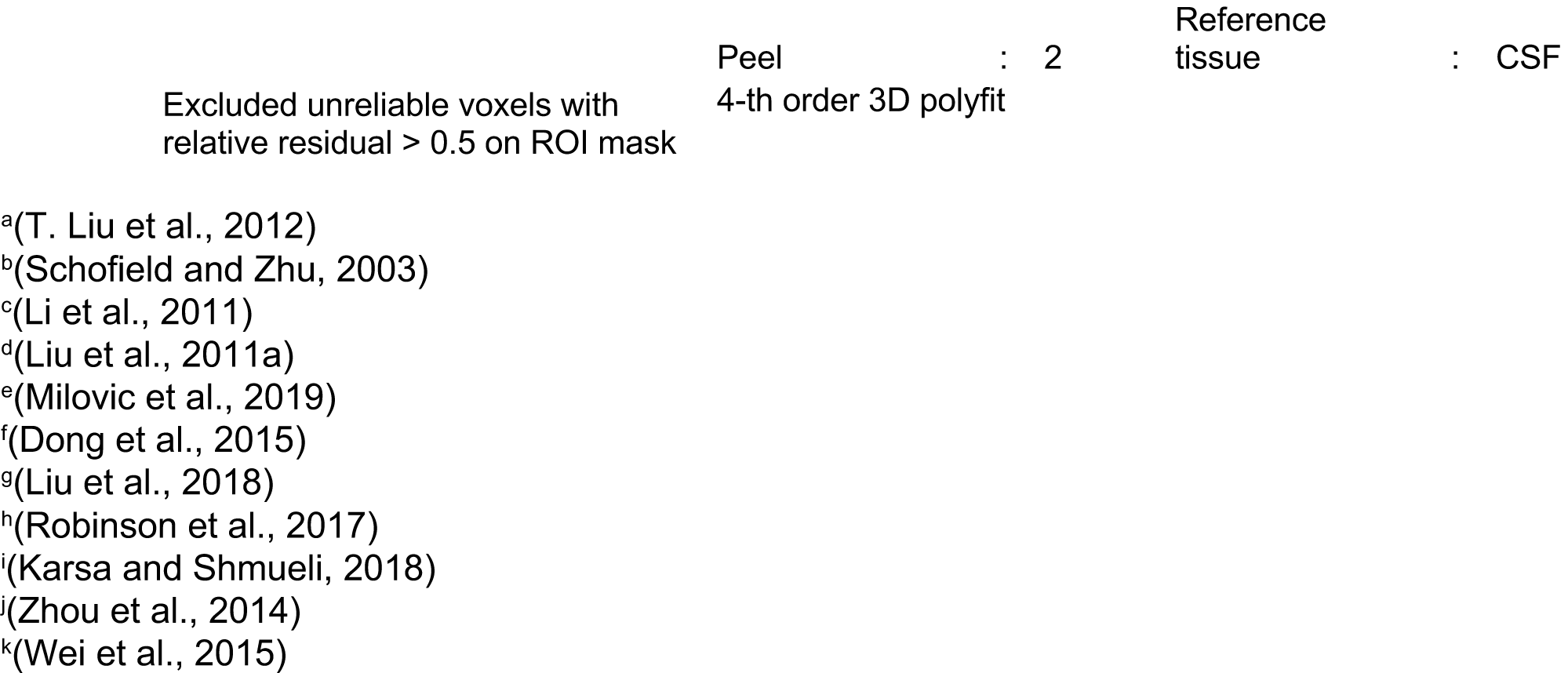
Methods and parameters used in the three QSM processing pipelines on the test data.

## 3. Results

Fig. 5 shows the resulting total field map, local field map and susceptibility map of each QSM processing pipeline to demonstrate the flexibility of mixing-and-matching methods from various toolboxes (indicated by the text colours). The total field map using the graph-cut approach is nearly identical to that of SEGUE, as the two algorithms were trying to unwrap the phase data by adding or subtracting a correct multiple of 2π to the wrapped phase data. The total field map generated by the Laplacian approach shows different brightness compared to the other two because some of the harmonic field components in the data were removed by the Laplacian operator as it has been previously described in the literature (Robinson et al., 2017; Schweser et al., 2013). The contrast of local field maps between all the three methods is very similar, whereas differences can be observed mainly on the edges of the brain. All three pipelines produced high quality susceptibility maps from the same input data. Notably, the distinct approaches to incorporate prior information in the dipole inversion optimisation created slightly different appearances between the resulting susceptibility maps, where FANSI with weak-harmonic regularisation and MEDI+0 gave the smooth results, while the result of Star-QSM looks more realistic using the selected algorithm parameters.

**Fig. 5:**
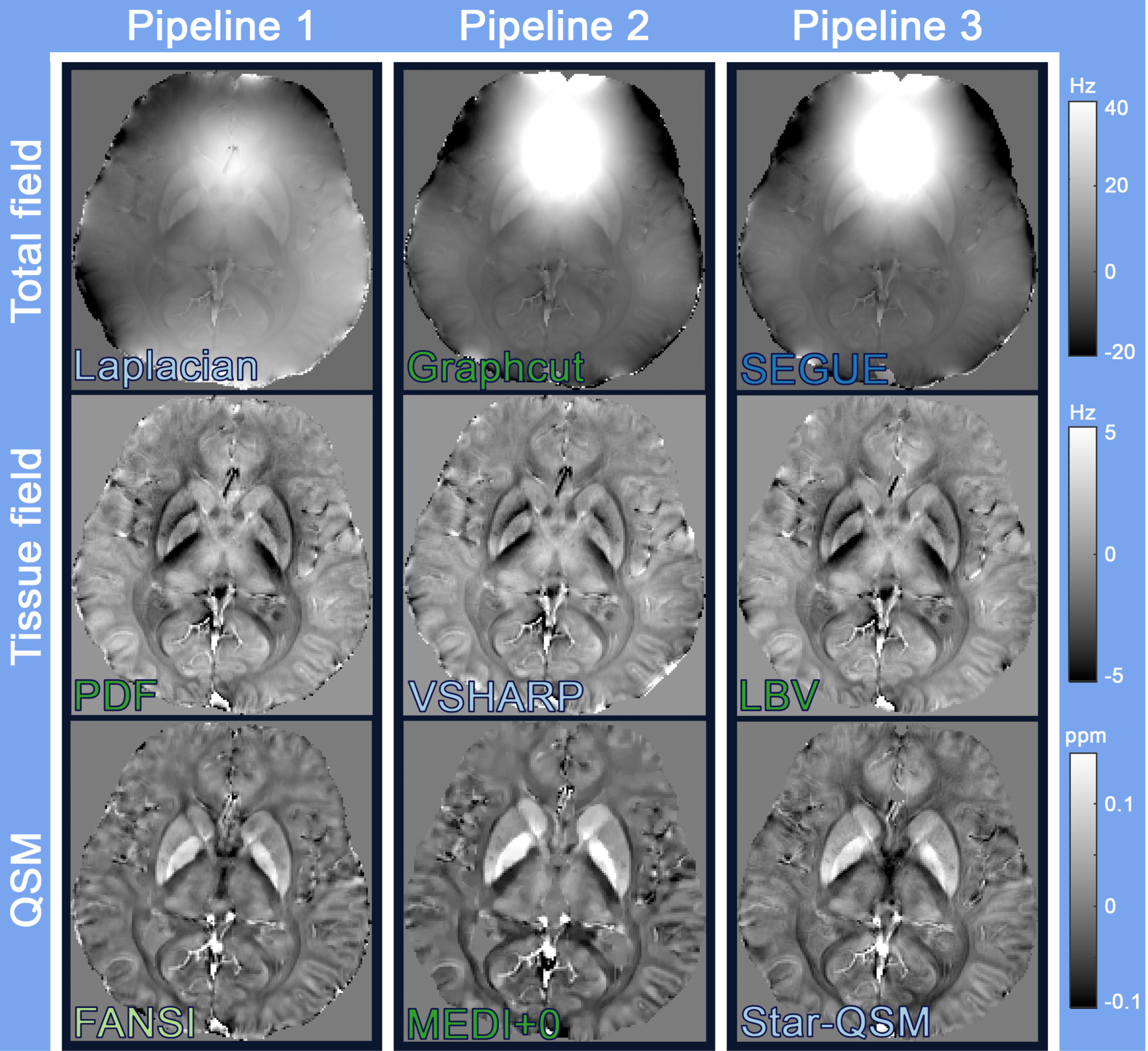
QSM processing results with SEPIA. The same *in vivo* data was processed in three different QSM processing pipelines with the combination of methods from various toolboxes. The susceptibility maps obtained from the three pipelines show similar contrast, particularly between the deep grey matter structures where the highest susceptibility values in globus pallidus reflecting its high iron deposition. The quality of the results is dependent on a variety of factors such as the kind of prior information in the regularised optimisation, regularisation weighting and the quality of the input tissue field map. For studies included QSM, we suggest using a single pipeline for all subjects to minimise the variation introduced by different methods. Colour of the method representing toolbox: light blue - STI Suite; blue - SEGUE; light green - FANSI toolbox; green - MEDI toolbox.

Fig. 6 shows the high-pass filtered phase at the last echo on 1 slice together with the magnitude, SWI and SMWI images at the last echo with minimum intensity projection on an 8-mm slab. The positive phase SWI image (pSWI, Fig. 6d) shows the most extensive venous structures compared to the original magnitude (Fig. 6b) and paramagnetic SMWI images (pSMWI, Fig. 6f), benefiting from the effect of signal phase extends beyond the structure itself, also known as blooming artefact. However, the contrast in the pSWI image is also affected by the enhancement from the residual non-local fields which are more persistent toward the tissue/bone or air sources such as frontal sinus (green arrows). Since SMWI used a QSM map which has undergone a more sophisticated background field removal, these effects are not an issue for pSMWI, as all non-local field contributions were removed in QSM processing. Improved contrast localisation can also be observed on the diamagnetic SMWI image (dSMWI, Fig. 6e). It is clear that the dSMWI image enhances the optic radiation and forceps major in white matter (orange arrows), as is also the case on the negative phase SWI image (nSWI, Fig. 6c). Yet, in the nSWI, this contrast enhancement comes at the cost of artefacts surrounding the putamen and the globus pallidus (purple arrows), where the negative in-plane field surrounding the structures (see Fig. 6a and tissue field in Fig. 5) creates a blooming artefact that both reduces the edge definition between the structures and undesirably extends the external capsule contrast farther beyond the fibre itself.

**Fig. 6:**
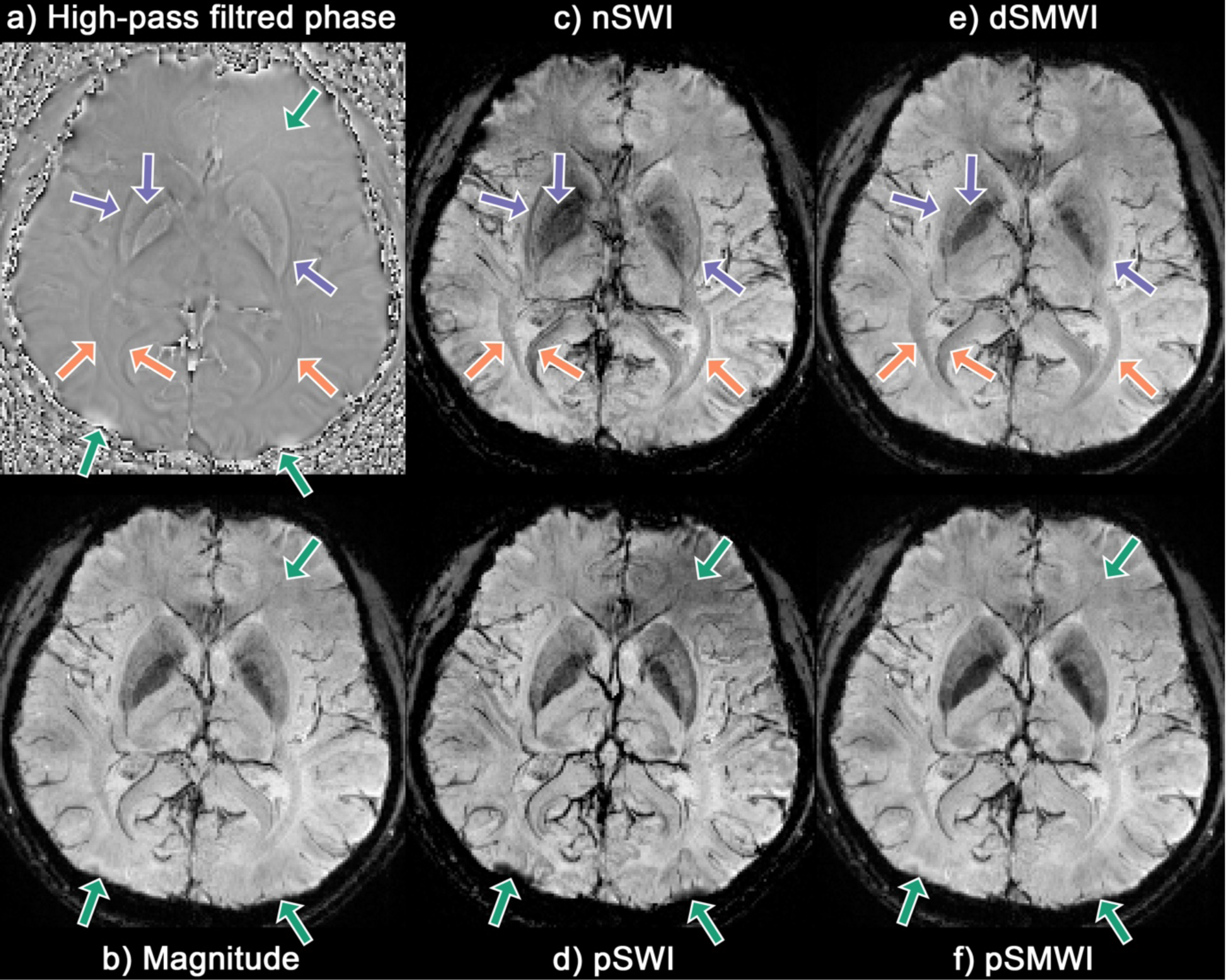
(a) High-pass filtered phase image of 1 slice (display range=[-π,π]), minimum intensity projection images on an 8-mm slab of the (b) magnitude GRE, (c) negative phase SWI (nSWI), (d) positive phase SWI (pSWI), (e) diamagnetic SMWI (dSMWI), and (f) paramagnetic SMWI (pSMWI) at the last echo (TE=45ms). The venous structures are more pronounced in the pSWI image compared to the magnitude and pSMWI results. Green arrows indicate the regions affected by non-local fields contrast enhancement in pSWI but not in pSMWI and original data. Purple arrows indicate that the blooming artefact originated from the globus pallidus and the putamen makes the two structures less distinguishable and creates a misleading contrast in the outer part of the putamen similar to the external capsule. Orange arrows indicate the white matter fibre bundle enhancement on optic radiation and the splenium of the corpus callosum using the negative phase/susceptibility image.

## 4. Discussion

SEPIA is a pipeline tool dedicated to QSM processing to deliver a connection between various QSM software packages in a single platform. By doing so, users can effortlessly experiment with different method combinations in SEPIA. Unlike other quantitative imaging techniques such as T_1_ mapping or T_2_* mapping where a standardised protocol and processing method can be deployed, the ill-posed nature of QSM hinders such standardisation and the results are often depended on the data quality such as SNR and image artefacts. Therefore, extending QSM processing flexibility by expanding the number of available options can be crucial to achieving high quality QSM results.

The user-friendly GUI of SEPIA presents an intuitive way to construct a processing pipeline without the requirement of users to have advanced programming skills. All supported methods are shown as options with the corresponding adjustable parameters presented in the GUI. This provides easy access to all the possibilities available in SEPIA to construct a pipeline. Together with other utility functions (see sections 2.3.2-2.3.5) which can be helpful to further improve QSM results, it is particularly useful for researchers who want to include QSM in their studies but with less experience on the processing and theory.

The main objective of SEPIA is to provide greater flexibility in QSM research, hence the ability to incorporate new methods is also its major focus. Through the wrapper function design in the communication level, developers can embed their methods in the existing SEPIA framework independently and without any updates on their codes or regularisation parameter (value often depended on input data unit), because all the input data is accessible in the wrapper function and they can be adjusted before and after the main process, which is time-efficient in terms of method development. Developers are also encouraged to provide their methods and wrappers to the SEPIA development team, such that the methods can be officially supported as add-ons in SEPIA future releases. This can bring a mutual benefit to both users and developers, where SEPIA serves as a platform for developers to share their methods and users can utilise the most advanced techniques for their research.

For readers who have less experience with QSM processing and SEPIA, a step-by-step tutorial is provided in the documentation website (https://sepia-documentation.readthedocs.io/) to demonstrate how to construct their first pipeline in SEPIA. However, given a vast number of method combinations available in SEPIA, beginners could find it challenging on how to optimise a pipeline for their data. Here, we provide some practical considerations for choosing phase processing methods. For total field recovery and phase unwrapping, path-following methods such as SEGUE are generally considered more accurate, while Laplacian-based approaches are more robust to noise (Robinson et al., 2017). We, therefore, suggest users start with path-following methods before trying the faster Laplacian-based approaches. A recent review of background field removal methods demonstrated that tissue fields obtained by most methods have minimal differences especially in deep brain regions (Schweser et al., 2016b), while methods based on the assumption of no susceptibility sources close to region-of-interest boundaries, such as SHARP and V-SHARP, should be used with caution in studying cortical regions of the brain. A detailed summary of background field removal methods including benefits and limitations can be found in (Schweser et al., 2016b). The optimal method for dipole inversion highly depends on the needs of the researcher comprising factors such as data quality and tissue of interest of the study, trading data fidelity for data consistency (Bilgic et al., 2014b). The authors’ experience suggested that methods incorporating image-based information such as SNR map or magnitude images can produce susceptibility maps with less streaking artefact. Users should also investigate the optimal regularisation value for their data instead of relying on the default setting in SEPIA to avoid under- or over-regularised result.

The advance of QSM methodology in the last decade makes clinically acceptable susceptibility maps possible. However, the limitations of QSM should not be overlooked. Firstly, ensuring accurate intermediate results of the QSM processing is of uttermost importance to produce a high quality QSM map, as error can easily propagate from one step to another. This includes not only general phase processes but also pre-processing steps such as the definition of the region of interest (e.g. brain extraction). SEPIA can simplify the workflow of QSM processing, yet users should guarantee the quality of the results before proceeding to the next one. Secondly, for inter-subject comparison or comparison to literature studies, a reference tissue/region must be selected for referencing the susceptibility map in the subject level since QSM provides only a relative measure instead of absolute values (Deistung et al., 2016). Determination of an appropriate reference region is still an active investigation in QSM as factors such as inter-subject variability have to be carefully considered. Lastly, it is widely accepted that current QSM model, based on the deconvolution of a dipole field, holds in tissue with only isotropic susceptibility or randomly oriented structures (Liu, 2010; Yablonskiy and Sukstanskii, 2015) Thus, interpretation of susceptibility in white matter should be taken with extra caution. Anisotropic susceptibility has been previously demonstrated with myelinated axons (Lee et al., 2010; Li et al., 2012; Wharton and Bowtell, 2015), accounting for the fibre-orientation dependency in QSM with the current model. Additional phase variation in early echo times induced by white matter microstructure is another confounding factor of studying QSM (Gelderen et al., 2011; Sati et al., 2013; Wharton and Bowtell, 2013).

Future development of SEPIA will focus on simplification of adding a third-party method to the framework. Currently, adding a new method to SEPIA is entirely based on scripting, which may not be straightforward to perform. Having a GUI based operation to install a new method as an add-on can allow users to develop, exchange or utilise new methods in a simpler manner. In our current implementation, we support the traditional workflow used by most researchers, which separates background field removal from the QSM computation. Yet, single-step methods have also found some attention recently (Chatnuntawech et al., 2017; Liu et al., 2017; Sun et al., 2018). Although we do not currently support any of these methods, this would be straightforward to implement without any structural modification on the existing software architecture. Another focus will be to explore the possibility to incorporate processing methods that are not developed in MATLAB. Emerging techniques using deep learning have already shown very promising results with further improvement in QSM artefact reduction (Bollmann et al., 2019; Chen et al., 2019; Jung et al., 2020; Polak et al., 2020; Wei et al., 2019; Yoon et al., 2018; Zhang et al., 2020). As the implementations of these methods are primarily in Python, while there are rich resources already available in MATLAB and supported in SEPIA to perform tasks such as phase unwrapping and background field removal, it would be valuable for SEPIA to connect the two. Up to now, most deep learning methods do not yet have the flexibility to cope with arbitrary slice orientation and resolution (Jung et al., 2020; Yoon et al., 2018), this would therefore require data re-sampling to a pre-defined space which can be done with a plethora of software available for neuroimaging data, but we believe this is to be outside the scope of this toolbox. We also consider the compatibility between the SEPIA input and outputs data with BIDS (Gorgolewski et al., 2016), moving toward the standardisation of input and output phase data format and structure. Currently, if the user provides an output filename with a prefix (see Table 1) that is BIDS compatible, the resulting output is also naturally BIDS compatible. Automatically processing data with BIDS input would involve a further extension of the current BIDS specifications and therefore requires community effort before consensus is to be reached.

## 5. Conclusions

SEPIA is a pipeline toolbox for QSM processing with MRI phase data. It provides a connection between various QSM software packages in the field and a user-friendly GUI frontend for users to construct phase data processing pipelines without the requirements of advanced programming skills. The package is open-source and highly extendable, allowing developers to expand its capability by deploying new methods to the framework. The online documentation of SEPIA accommodates all essential information about QSM processing and SEPIA, vital for less-experienced researchers who are interested in including QSM in their research.

## Acknowledgement

This work was funded by the Netherlands Organisation for Scientific Research (NWO) with project number FOM-N-31/16PR1056.

## Notes

https://github.com/kschan0214/sepia

### Competing Interest Statement

The authors have declared no competing interest.

