## Supplementary A for "SEPIA - SuscEptibility mapping PIpeline tool for phAse images"

### Supplementary material A

#### Using directory as an input method in SEPIA

Table S1: Directory organisation to be used as SEPIA processing pipeline input.

| **Data** | **Identifier** | **Example** |
| --- | --- | --- |
| **One-step QSM processing** | | |
| Phase image | *ph* | sub-01_part-phase_gre.nii.gz |
| Magnitude image | *mag* | sub-01_part-mag_gre.nii.gz |
| Header | *header* | sepia_header.mat |
| Mask^†^ | *mask* | sub-01_gre_mask.nii.gz |
| **Phase unwrapping application** | | |
| Phase image | *ph* | sub-01_part-phase_gre.nii.gz |
| Magnitude image | *mag* | sub-01_part-mag_gre.nii.gz |
| Header | *header* | sepia_header.mat |
| Mask^†^ | *mask* | sub-01_gre_mask.nii.gz |
| **Background field removal application** | | |
| Total field map | *total-field* | sub-01_part-phase_gre.nii.gz |
| Header | *header* | sepia_header.mat |
| Mask | *mask* | sub-01_gre_mask.nii.gz |
| Noise SD map^††^ | *noise-sd* | Sepia_noise-sd.nii.gz |
| **QSM application** | | |
| Local field map | *local-field* | sub-01_part-phase_gre.nii.gz |
| Header | *header* | sepia_header.mat |
| Mask | *mask* | sub-01_gre_mask.nii.gz |
| Magnitude image^††^ | *mag* | sub-01_part-mag_gre.nii.gz |
| Weighting map^††^ | *weights* | Sepia_weights.nii.gz |
| ^†^Optional input data |  |  |
| ^††^Mandatory for some algorithms | | |

#### Mathematical formalisms used in SEPIA processing

##### Noise estimation in the phase data and SNR weighted image

The prior estimate of the standard deviation of noise in the phase data (noise-sd.nii.gz) using the non-linear temporal phase unwrapping method from MEDI toolbox is derived from:

$${noiseSD}_{MEDI}=\sqrt{\frac{\sum_{k=1}^{K} \left| M_{k} \right|^{2}}{\left( \sum_{k=1}^{K} \left| M_{k} \right|^{2} \right)\left( \sum_{k=1}^{K} \left| M_{k} \right|^{2}{t_{k}}^{2} \right)-\left( \sum_{k=1}^{K} \left| M_{k} \right|^{2}t_{k} \right)^{2}}} [Eq.S1a]$$

where $M_{k}$ and $t_{k}$ is the magnitude signal and echo time in second at echo k.

The prior estimate of the standard deviation of noise in the phase data using the optimum weight combination for temporal phase unwrapping is derived from

$${noiseSD}_{OW}=\sqrt{\frac{1}{\sum_{k=2}^{K} \frac{\left( t_{k}-t_{1} \right)^{2}M_{k}^{2}M_{1}^{2}}{M_{k}^{2}+M_{1}^{2}}}}=\sqrt{\frac{1}{\sum_{k=2}^{K} \frac{1}{\sigma_{\omega_{k1}}^{2}}}} [Eq.S1b]$$

To match the order of magnitude of the standard deviation derived from the two methods, the noise standard deviation estimated from the optimum weight method is further normalized by the Euclidean norm inside the signal mask.

$${noiseSD}_{OW,SEPIA}=\frac{{noiseSD}_{OW}}{{norm}_{Mask}({noiseSD}_{OW})} [Eq.1c]$$

Consequently, the SNR weighting map (weights.nii.gz) is computed by

$$weights=\left\{ \begin{aligned} 0, &noiseSD=0. \\ \frac{1/{noiseSD}}{{max}_{Mask}(1/{noiseSD})}, &otherwise. \end{aligned} \right. [Eq. S2]$$

which is the inverse standard deviation of noise in phase normalised by the maximum value in the resulting map.

##### Relative residual from mono-exponential model fitting with multi-echo data

Multi-echo complex-valued signal is simulated after the estimation of the unwrapped total field map *f*:

$$S_{simulated,k}=S_{0}e^{-R_{2}^{*}t_{k}}e^{i2\pi ft_{k}} [Eq. S3]$$

where $t_{k}$ is the time in second at echo *k*, $S_{0}$ and $R_{2}^{*}$ are estimated using the approach in (Khabipova et al., 2015). Subsequently, the relative residual is computed from

$$relative residual= \frac{\sum_{k=1}^{K} \left| \tilde{S}_{k}-\tilde{S}_{simulated,k} \right|}{\sum_{k=1}^{K} \left| \tilde{S}_{k} \right|} [Eq. S4a]$$

$$\tilde{S}_{k}= S_{k}\bar{S_{1}} [Eq. S4b]$$

$$\tilde{S}_{simulated, k}= S_{simulated,k}\bar{S}_{simulated,1} [Eq. S4c]$$

where $S_{k}$ is the measured complex-valued signal at echo k. The complex conjugate operation in Eq. S4b-c is essential to avoid the mismatch between the measured and simulated signal regarding the phase contribution from B1 in all echoes without explicit estimation of the effect and assuming that phase errors are minimal on the first echo.
